# CNS diseases cerebrospinal fluid single-cell atlas reveals immune characteristics of neuropsychiatric systemic lupus erythematosus

**DOI:** 10.64898/2026.03.31.715151

**Authors:** Xin-Jie Wang, Shang-Zhu Zhang, Si-Yuan Fan, Wen-Jia Zhang, Tian-Yi Ma, Wei-Ting Fang, Nan Liang, Yang Wu, Shi-Qi Yang, Chen-Rui Xia, Zi-Fan Zhao, Jiu-Liang Zhao, Dong Xu, Xiao-Feng Zeng, Hong-Zhi Guan, Yang Ding, Ge Gao, Meng-Tao Li

## Abstract

Neuropsychiatric systemic lupus erythematosus (NPSLE) is a potentially severe complication of systemic lupus erythematosus (SLE), yet its pathogenesis remains largely elusive. By jointly probing the immune dynamics of subjects’ cerebrospinal fluid (CSF) and peripheral blood, we showed that both innate and adaptive immune responses jointly contribute to the pathogenesis of NPSLE. In particular, we found the remarkable enrichment of BAM-CCL3, a subtype of border-associated macrophages with strong recruitment capacity, implicating its potential role in central nervous system (CNS) inflammation. We also observed pronounced activation of memory B cells and CD4^+^ regulatory T cells in NPSLE CSF, along with the preferential blood-to-CSF migration and subsequent within-CSF clonal expansion of CD8^+^ effector memory T cells in NPSLE patients, suggesting a persistent CNS-localized adaptive immune dysregulation. Finally, we developed the single-cell CNS disease CSF–Blood Atlas (scCDCB), a comprehensive collection for CSF and peripheral blood of multiple CNS diseases, which is publicly available at (https://sccdcb.gao-lab.org) to serve as a reference for future research on CNS diseases.

## Introduction

Neuropsychiatric systemic lupus erythematosus (NPSLE) is a potentially severe complication of systemic lupus erythematosus (SLE) [**1**]. Approximately 50% of SLE patients develop neuropsychiatric manifestations during the disease course, with one third attributable to the lupus erythematosus, leading to greater organ damage, functional disability, and a higher risk of mortality[2,3].

Given the intrinsic difficulty in obtaining brain tissue *in vivo*, investigating NPSLE’s pathogenic mechanisms is particularly challenging and CSF essentially represents a critical window for probing the cellular and molecular dynamics driving NPSLE and other CNS-related disorders [4–14].Here we curated the largest single-cell CNS diseases atlas, covering 1.13 millions cells derived from CSF and paired peripheral blood samples for 8 central nervous system (CNS) disorders. Empowered by the first reported single-cell CSF dataset for NPSLE collected at Peking Union Medical College Hospital (PUMCH), we characterized the global dynamics for NPSLE’s immune microenvironment, including a pronounced pro-inflammatory tendency and activation of various immune cells, highlighting the joint contribution of both innate and adaptive immune responses to the pathogenesis of NPSLE. Moreover, we also showed the systematic difference between CSF and peripheral blood in term of cell composition, demonstrating at the PBMC samples should not be taken as a proxy for CSF in the study of CNS-related disorders. The whole atlas is available publicly as an online database single-cell CNS disease CSF–Blood Atlas (scCDCB) at https://sccdcb.gao-lab.org, aiming to a reference for follow-up basic and clinical studies for the community.

## Results

We gathered 19 studies from published datasets, including 228 CSF samples and 64 blood samples (Table 1 and Table S1, Fig. 1A). In addition, we collected CSF samples from 11 patients with SLE at PUMCH. Among them, nine patients had NPSLE during active disease episodes (Fig. 1A) and were referred to as the NPSLE group in subsequent analyses. In the sixth patient (Table S2), serial CSF samples were obtained at three time points: before treatment, after glucocorticoid pulse therapy and prior to the first intrathecal injection, and at the time of the second intrathecal injection. The rest two patients did not suffer from NPSLE during active disease episodes upon sampling and, while used in data integration, were not included into the NPSLE group above; in particular, one patient had a history of NPSLE, but the sample was collected when the neuropsychiatric symptoms were not active, and one patient had SLE without CNS involvement. In total, 13 paired CSF and blood samples were collected. After strict quality control and batch effect correction(Fig. S1), a total of 1,131,445 cells were retained, composed of 601,239 PBMCs and 530,206 CSF cells, and subsequent integration produced 82 cell types (Fig. 1B, C).

**Fig. 1.**
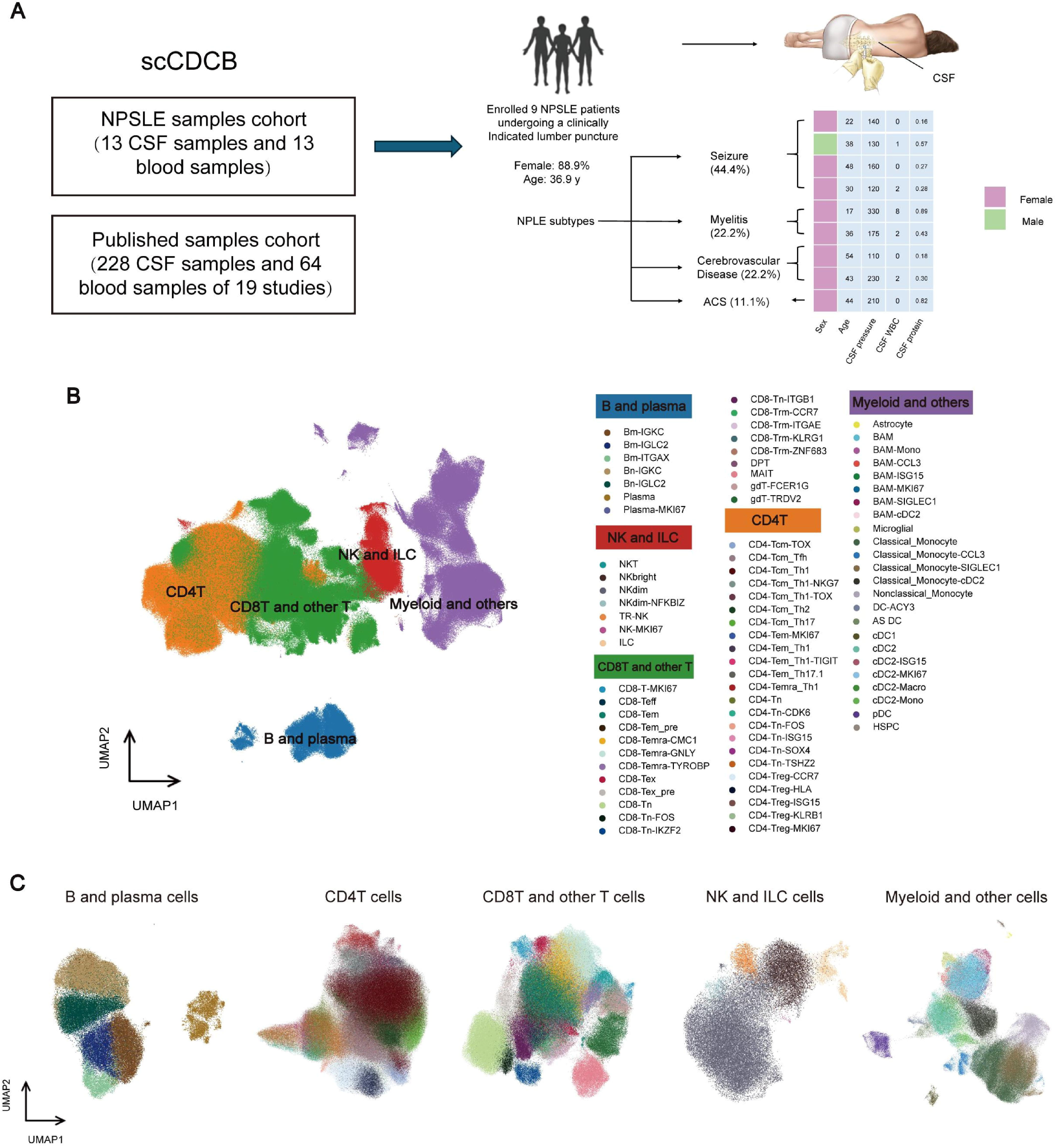
Single-cell CNS disease CSF – Blood Atlas and immune microenvironment composition. **A.** Establishment of the NPSLE samples cohort. Units for CSF opening pressure (mmH₂ O), white blood cell count (cells/µL), and protein concentration (g/L) are shown. **B.** UMAP plot showing the major cell types, colours represent different cell populations. **C.** UMAP plot showing the subtypes, To facilitate illustration, cells are grouped into five panels: B cells, CD4^+^ T (CD4T) cells, CD8^+^ T (CD8T) cells, NK cells, and myeloid cells.

**Table 1.** Samples cohort.

| **Dataset** | **Disease** | **CSF**<br>**samples** | **CSF**<br>**VDJ**<br>**samples** | **Blood**<br>**sample**<br>**s** | **Blood VDJ**<br>**samples** |
| --- | --- | --- | --- | --- | --- |
| E-MTAB-9916<sup>[38]</sup> | HC | 3 | / | / | / |
| GSE241385<sup>[39]</sup> | HC | 12 | / | / | / |
| GSE243905<sup>[40]</sup> | HC | 3 | 3 | 6 | 6 |
| GSE252701<sup>[41]</sup> | / | 33 | / | / | / |
| GSE133028<sup>[13]</sup> | MS,HC | 12, 3 | / | 11, 3 | / |
| GSE134577<sup>[14]</sup> | HC,AD,MCI | 6, 3, 3 | 6, 3, 3 | / | / |
| GSE138266<sup>[42]</sup> | MS,IIH | 6, 6 | / | 5, 5 | / |
| GSE141578<sup>[12]</sup> | PD,HC,DLB | 9, 2, 2 | / | / | / |
| GSE163005<sup>[43]</sup> | IIH,MS | 5, 5 | / | / | / |
| PRJNA717310<sup>[44]</sup> | HC | 3 | 3 | / | / |
| GSE172003<sup>[45]</sup> | MS | 6 | / | / | / |
| GSE194078<sup>[46]</sup> | HC,IDD,MS | 2, 6, 3 | / | 2, 6, 3 | / |
| GSE200164<sup>[47]</sup> | HC,MCI/AD | 45, 14 | / | / | / |
| GSE202410<sup>[48]</sup> | HC | 4 | / | 4 | / |
| PRJNA866296<sup>[49]</sup> | MS | 6 | / | / | / |
| GSE232391<sup>[50]</sup> | SMA | 22 | / | / | / |
| GSE233917<sup>[51]</sup> | MS | 2 | / | / | / |
| GSE234069<sup>[52]</sup> | IIH | 2 | 2 | / | / |
| NPSLE | NPSLE,SLE<br>without central<br>nervous system<br>involvement,NPSLE<br>in sustained<br>remission | 11,1,1 | 10,1,1 | 11,1,1 | 11,1,1 |
| HRA003831<sup>[53]</sup> | SLE,HC | / | / | 15, 4 | 15, 4 |

### Both Cell Compositions and Cell States Changed in NPSLE

We extracted CSF samples and performed immune microenvironment analyses using the same analytical framework. The immune microenvironmental composition of NPSLE CSF differs markedly from that of other CNS diseases or HCs (Fig. 2A). Notably, many cell types exhibited significantly increased proportions compared with HCs, even after adjusting for age and sex as covariates (Fig. 2B). In addition, most these proportionally differential cell types also exhibit a large number of upregulated genes in NPSLE CSF compared with HC CSF (Fig. 2C; Table S3), indicating that NPSLE CSF undergoes a pronounced shift not only in the relative abundance of specific cell types but also in the cellular states within each cell type. Of note, the majority of differentially expressed genes appeared to be upregulated in a cell type-specific manner (Fig. 2D), suggesting distinct modes of state alterations across different cell types, which we examine in detail below.

**Fig. 2.**
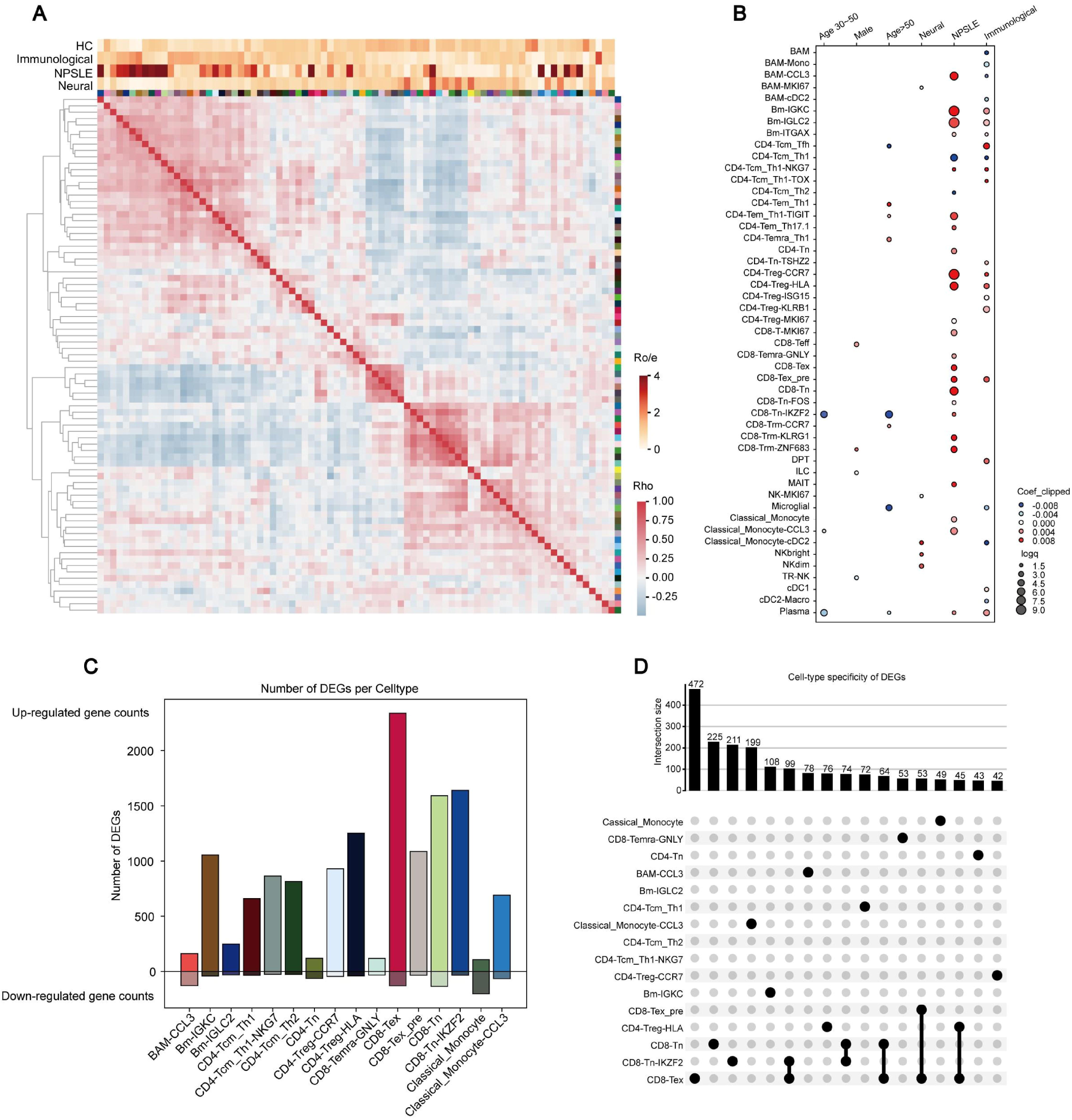
Composition of the immune microenvironment in NPSLE CSF. **A.** The heatmap of the Spearman’s correlation coefficients (ρ) on cell type proportions between different CSF patients, with the top four bars shows the distribution of the ratio of enrichment (Ro/e) for each cell type across different diseases (or HC). Cell type colors follow those in Fig. 1C. **B.** (GLM regression based on CSF cells, jointly considering disease, age, and sex, with FDR < 0.05. **C.** The numbers of those up- and down-regulated DEGs in each cell type filtered with |logFC| >1 and FDR<0.05. Only those cell types with more than 100 such upregulated DEGs are shown. **D.** The distribution of upregulated DEGs between different cell types. Only those patterns of upregulation across cell types with more than 40 matching DEGs are shown.

### BAM-CCL3 Recruits Immune Cells in NPSLE and Is Associated with Pro-inflammatory and Tissue-damaging Responses

We first focused on the myeloid lineage and found the BAM-CCL3, a subtype of BAMs that has not been previously reported, yet displaying a strong proportional increase in NPSLE (Fig. 2B). Compared to other myeloid cells, BAM-CCL3 upregulates chemokine genes such as *CCL3*, *CCL4*, and *CXCL8*, and also the pro-inflammatory cytokine *IL1B* [15], suggesting a strong capacity of recruiting immune cells and promoting inflammatory response in the CNS (Fig. 3A). Compared to HCs, the transcription profile of NPSLE BAM-CCL3 mainly reflects an even stronger pro-inflammatory capabilities (Fig. 3B, C, Fig. S3A). This includes further upregulation of many chemokine genes, as well as genes related to adhesion capacity regulation, macrophage phagocytosis, and polarization, such as *LGMN* [16], *ABL2* [17], *NINJ1* [18], and *CXCL8* [19]. Interestingly, they also upregulate VEGF family genes, including *VEGFA* and *VEGFB*, which can promote endothelial proliferation and activation [20]; activated endothelial cells are more likely to recruit immune cells [21], potentially increasing BBB permeability. Such strong regulation of chemokine and pro-inflammatory genes compelled us to examine further whether BAM-CCL3 could recruit multiple immune subsets altogether, and cell-cell communication analyses supported this hypothesis, demonstrating that BAM-CCL3 recruits distinct immune cell subsets through distinct chemokine–receptor axes. Specifically, it attracts T cells via the CXCL16–CXCR6, CCL4–CCR5, and CCL3–CCR5 axes; myeloid cells through the CCL3L3/CCL3–CCR1 axis; and B cells, dendritic cells, as well as a subset of T cells via the CCL20–CCR6 axis (Fig. 3D). Overall, these results suggest a potentially indispensable role of BAM-CCL3 in promoting inflammation in patients with NPSLE.

**Fig. 3.**
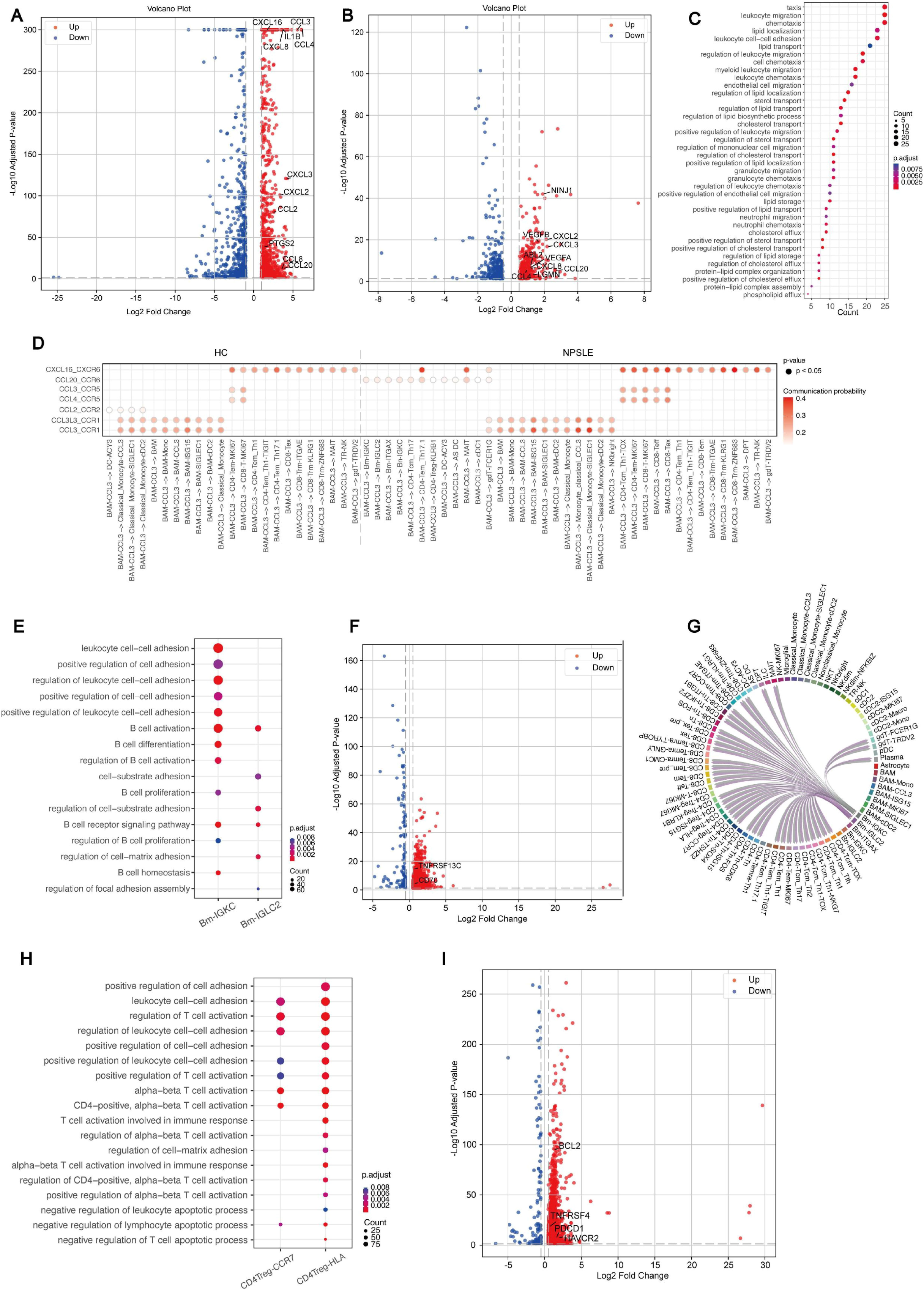
DEG, Pathways and Interactions Enriched by BAM-CCL3, Bm-IGKC, Bm-IGLC2, CD4Treg-CCR7 and CD4Treg-HLA in NPSLE CSF. **A.** DEGs of BAM-CCL3 relative to other myeloid cells in all CSF samples, filtered by |logFC| > 1 and FDR < 0.05. **B.** DEGs of BAM-CCL3 in NPSLE CSF relative to HC CSF, filtered by |logFC| > 0.5 and FDR < 0.05. **C.** The enriched pathways of the upregulated DEGs of BAM-CCL3 in Fig. 3B. Only pathways associated with leukocyte recruitment, chemotaxis, migration and/or lipids are shown. **D. D**istribution of communication probabilities of axes involving different CCL and CXCL chemokines sent out by BAM-CCL3, filtered by p<0.05. **E.** The enriched pathways using the upregulated DEGs of Bm-IGKC and Bm-IGLC2 in NPSLE CSF relative to HC CSF. Only pathways associated with B cell or cell adhesion were shown. **F.** DEGs of Bm-IGKC and Bm-IGLC2 in NPSLE CSF relative to HC CSF, filtered by |logFC| > 0.5 and FDR < 0.05. **G. Cell types possibly regulated by the** CD70-CD27 signaling pathway send out by Bm-IGKC and Bm-IGLC2 in NPSLE. The inner bars indicate the targets that receive signals from the corresponding outer segments. The size of each inner bar is proportional to the strength of the incoming signal received by that target. **H.** The enriched pathways using the upregulated DEGs of CD4Treg-CCR7 and CD4Treg-HLA in NPSLE relative to HC. Only pathways associated with T cell activation, cell adhesion and/or apoptotic process were shown. **I.** DEGs of CD4Treg-CCR7 and CD4Treg-HLA in NPSLE CSF relative to HC CSF, filtered by |logFC| > 0.5 and FDR < 0.05.

### Bm-IGLC2 and Bm-IGKC are Activated in NPSLE and Probably Activate Other Immune Cells via the CD70-CD27 Pathway

We then moved to subtypes of B cells that are known to play a major role in autoimmune diseases. In NPSLE, Bm-IGLC2 and Bm-IGKC, two subsets of memory B cells proportionally increased in NPSLE patients (Fig. 2B), upregulate immune response activation pathways relative to HCs, indicating an activated state (Fig. 3E). This activation may be due to BAFF and APRIL, two key players in B cell activation[22], as the information flow of BAFF and APRIL with Bm-IGLC2 and Bm-IGKC in NPSLE is stronger than that in HC, and memory B cells in NPSLE upregulate expression of *TNFRSF13C* (Fig. S3B, Fig. 3F); this is also consistent with previous findings of elevated BAFF and APRIL in the CSF of SLE and NPSLE patients[23]. Additionally, such activated memory B cells in NPSLE further upregulate its expression of *CD70* (Fig. 3F), consistent with previous studies on antigen- encounter-activated B cells[24], and utilize the CD70-CD27 axis to regulate other immune cells (Fig. S3C), including T and B cells (Fig. 3G, Fig. S3D), possibly affecting their survival and activation, forming more memory B and plasma cells[24] and ultimately exacerbating CNS inflammation.

### Long-lived CD4Treg-CCR7 and CD4Treg-HLA Cells Exert Potent Immunosuppression in NPSLE

We then shifted our focus to T cells, another key component in adaptive immunity and autoimmune diseases. We found that CD4Treg-CCR7 and CD4Treg-HLA, two subsets of CD4Treg cells proportionally increased in NPSLE (Fig. 2B), are also activated and display an anti-apoptotic signature in NPSLE compared to HCs (Fig. 3G). Because Treg cells are known to mainly exert immunosuppressive capabilities [25], such strong activation suggests immunosuppressive activity of Treg cells within the NPSLE CSF microenvironment, possibly as an immune compensatory response against the pathological overactivation of other immune cells. Indeed, cell-cell communication analyses revealed that, compared with HCs, CD4Treg-CCR7 and CD4Treg-HLA in NPSLE are likely to be more activated by CD70 (Fig. S3E) via the CD70-CD27 axis from memory B cells (Fig. 3F and Fig. S3D). In addition, CD4Treg-CCR7 and CD4Treg-HLA may also be activated through other interactions, such as the CD86-CD28 axis associated with the survival and activation of CD4Treg cells (Fig. S3E, Fig. S3F). Functionally, these CD4Treg cells may utilize the CD86-CTLA4 axis to suppress antigen-presenting cells’ (CD86-CD28-axis-based) co-stimulation of other T cells, by inducing their trans-endocytosis of CD86 surface proteins (Fig. S3E, Fig. S3F) [26]. Moreover, these activated CD4Treg cells in NPSLE also upregulate co-inhibitory receptors such as *PDCD1* and *HAVCR2*, in addition to anti-apoptosis-related genes such as *BCL2* and *TNFRSF4* (Fig. 3I), potentially further enhancing their survival and immunosuppressive capabilities.

### CD8-Tem exhibits intense clonal expansion in the NPSLE CSF and possesses strong tissue migration capabilities

Finally, we used single-cell TCR sequencing data to identify characteristic clonal patterns of T cells in NPSLE CSF. We found that, despite an overall markedly lower clonal expansion in CSF T cells compared to blood T cells (Fig. S4A-C), clonally expanded T cells in NPSLE CSF still display a strong enrichment in CD8Tem cells whereas no particular enrichment was observed for HC CSF T cells, and also distinguish themselves from blood-expanded T cells that are mostly CD8Temra-biased (Fig. 4A, Fig. S4D-G). In addition, among the approximately 20–30% shared expanded clones between paired blood and CSF of NPSLE patients (Fig. 4B), the major T subtypes switch from mostly CD8Tem and CD8Temra in blood to mostly CD8Tem only in CSF (Fig. 4C), suggesting that CSF in NPSLE patients maintain a pool of CD8Tem cells from the blood (at least partially) to exert certain functions, possibly related to NPSLE pathology. In line with this, NPSLE CD8Tem cells exhibited the strongest blood-CSF migratory capacity among T subtypes (Fig. 4D), and their CSF part upregulates effector-like pathways associated with antigen response and cytotoxicity (Fig. 4E).

**Fig. 4.**
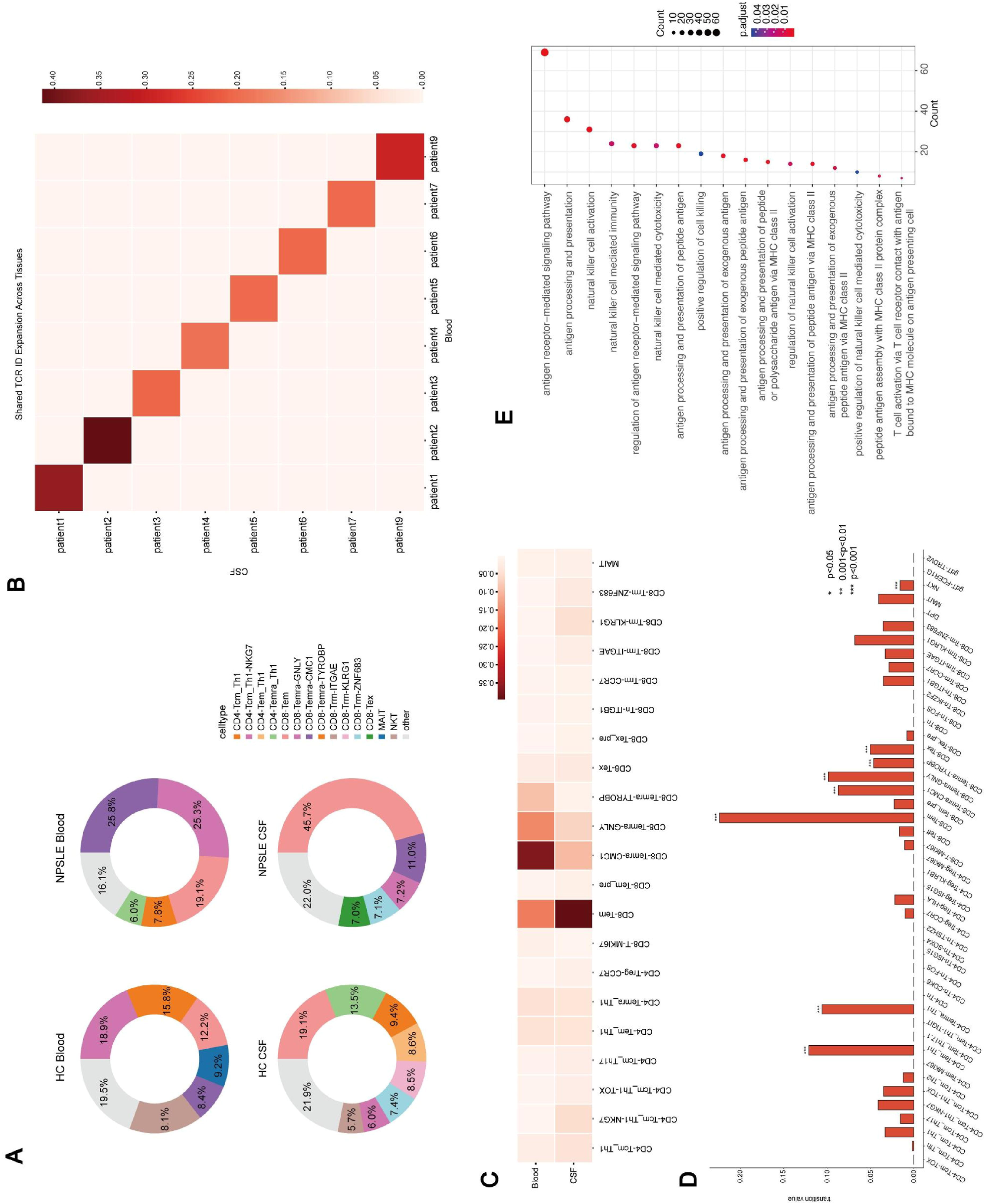
Clonal Expansion, Migration, and Differentiation of T Cells in CSF cells and PBMCs. **A.** The ratio of each cell type in the clonally expanded T cells in CSF and blood of NPSLE and HC patients. Clone IDs with more than five cells were selected as clonally expanded, and the proportion of each cell type within these expanded clone IDs was calculated. Cell types proportion less than 5% were grouped as “other”. **B.** The proportion of shared clone IDs from CSF cells and PBMCs in NPSLE patients. **C.** The distribution of shared clone IDs across cell types, with values representing the proportion of cells with shared clone IDs in CSF cells and PBMCs relative to the total cells of the tissue. **D.** STARTRAC-based migratory capacities of different cell types across NPSLE CSF and blood. **E.** The enriched pathways using the upregulated DEGs of CD8Tem in NPSLE relative to HC. Only pathways associated with antigen response and cytotoxicity were shown.

### scCDCB, the online portal for analyzing immune microenvironment of CNS diseases

To facilitate researchers in exploring the immune microenvironment of CNS diseases and their potential clinical applications, we provided the online scCDCB database based on the scCDCB atlas (http://47.95.3.103:8081/#/) (Fig. 5A). scCDCB database is continuously updated single cell RNA-seq open-source database. Users can access the scCDCB database through the Datasets dimension or the CellTypes dimension (Fig. 5B); specifically, the Datasets dimension enables the user to examine the atlas on a dataset basis, with the dataset ID, source link and cell type composition for each dataset displayed in its corresponding abstract block, while the CellTypes dimension groups cells by their cell types and lists for each cell type the total number of cells, clinical status and tissue types where these cells were discovered, as well as per-dataset description of this information in its abstract block. Users can further browse dataset details in the Datasets dimension, such as dataset metadata (Fig. 5C), cell composition (Fig. 5D), gene detection profile (Fig. 5E) and customized analyses (Fig. 5F). Users can also download the datasets of interest through our application programming interface (API) (Fig. 5G) for subsequent large-scale offline analyses.

**Fig. 5.**
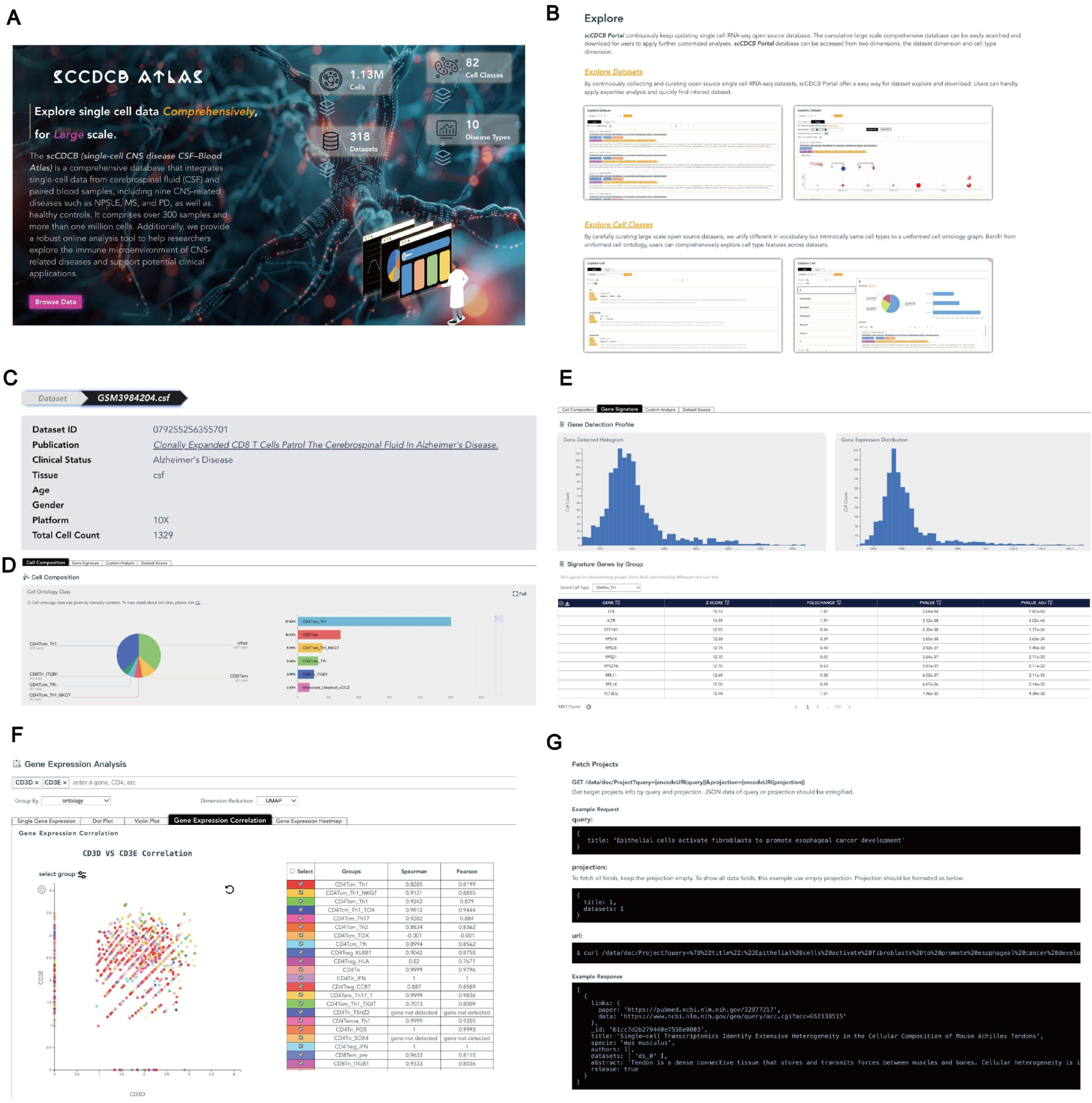
scCDCB database for future research of immune microenvironment of CNS diseases. A. The scCDCB database. B. the Datasets dimension and CellTypes dimension. C-F. Using the dataset GSM3984204 as an example, scCDCB database enables the users to examine its metadata (C), cell composition (D), distribution of the number of detected genes and total gene expression counts, as well as gene signatures (E), and various gene expression analyses, one of which shown here is the gene expression correlation analysis of two genes of interest (F). G. Users can also download the data using the API.

## Methods

### Patient Cohort

This study prospectively collected blood and CS<u>F</u> samples from patients with CNS NPSLE, as well as a limited number of patients with SLE without NP involvement, at PUMCH (Fig. 1A). All patients fulfill the 2019 European League Against Rheumatism/American College of Rheumatology (EULAR/ACR) classification criteria for SLE [27]. Neuropsychiatric involvement was diagnosed according to the 1999 ACR nomenclature and case definitions for NPSLE [28]. Patients with secondary neuropsychiatric manifestations, such as CNS infections or non – SLE-related CNS disorders, were excluded. The study cohort comprised patients with active NPSLE at disease onset, as well as one patient with NPSLE in sustained remission and one patient with SLE who experienced episodes of confusional arousals but did not meet the diagnostic criteria for NPSLE. For patients with active NPSLE, samples were obtained at the onset of NP symptoms, including seizures, cerebrovascular disease, myelitis, and acute confusional state (Table S2). All collected samples were incorporated into the construction of the scCDCB. For comparative analyses between NPSLE and HCs, only samples from patients with active NPSLE were included, unless otherwise specified.

Clinical data collected included demographics, clinical phenotypes of NPSLE, involvement of other organs, SLE Disease Activity Index (SLEDAI) score, C3 and C4 complement levels, autoantibody status, treatment regimens, and longitudinal follow-up information (Table S2). In addition, CSF parameters and neuroimaging findings were systematically recorded. CSF was obtained by lumbar puncture as part of routine clinical care. The study was approved by the Institutional Review Board of Peking Union Medical College Hospital (approval number I-23PJ671), and written informed consent was obtained from all participants.

### Single-Cell RNA Sequencing

The protoplast suspension was loaded into Chromium microfluidic chips with 30 (v2) chemistry and barcoded with a 10x Chromium Controller (10x Genomics). RNA from the barcoded cells was subsequently reverse-transcribed, and sequencing libraries were constructed with reagents from a Chromium Single Cell 30 v2(v2) reagent kit (10<u>x</u> Genomics) according to the manufacturer’s instructions. Sequencing was performed with Illumina (NovaSeq) according to the manufacturer’s instructions.

### Collection and Processing of CSF cells and PBMCs scRNA-seq Data

Single-cell RNA sequencing (scRNA-seq) data from CSF and blood were retrieved from published data in the form of SRA or BAM files. SRA files are converted to FASTQ format using fastq-dump from SRA Toolkit (version 2.9.6) with the parameter --split-files, and BAM files were converted to FASTQ format using cellranger bamtofastq. For samples without VDJ sequencing, the generated FASTQ files were processed with cellranger count. For samples with paired VDJ sequencing, data were processed using cellranger multi. All processing steps used 10x Genomics Cell Ranger v8.0.0[29], with the refdata-gex-GRCh38-2024-A reference genome and (for VDJ annotation only) the refdata-cellranger-vdj-GRCh38-alts-ensembl-7.1.0 reference.

### Quality Control

For CSF single-cell RNA-seq data, the Cell Ranger output h5ad files of each sample were loaded into SCANPY (v1.9.6) for downstream analyses including filtering, normalization, highly variable gene (HVG) selection, dimensionality reduction, and clustering. Ambient RNA contamination was estimated using SoupX [30] and corrected raw counts were subsequently used for downstream analyses. The percentage of counts in mitochondrial genes was calculated with scanpy.pp.calculate_qc_metrics. CSF cells were then filtered for those with 700–6000 detected genes and less than 10% counts in mitochondrial genes, and PBMCs were filtered for those with 700–6000 detected genes and less than 8% counts in mitochondrial genes. Potential doublets were identified with Scrublet[31] and removed if predicted_doublet=True or doublet_score > 0.3. Data normalization was performed using scanpy.pp. normalize_total, scaling the total counts per cell to 10,000, followed by log-transformation with scanpy.pp.log1p.

### Data integration and cell type annotation

Highly variable genes were selected using scanpy.pp.highly_variable_genes with batch_key set to “patient” and “tissue”. Batch effect correction was performed using scVI[32] with the random seed set to 0, batch_key specified as “patient” and “tissue”, two hidden layers, and a latent space dimensionality of 30. The scVI embeddings were then used for computing cell neighborhoods using scanpy.pp.neighbors with k=15, followed by UMAP computation with scanpy.tl.umap and clustering with scanpy.tl.leiden. Gene set scores were calculated using scanpy.tl.score_genes. Cluster-specific marker genes were identified using sc.tl.rank_genes_groups with the “wilcoxon” method, and cell type annotation was conducted based on canonical markers. Five major lineages were first annotated, including CD4^+^ T cells, CD8^+^ T cells, NK cells, B cells, and myeloid cells. Each major lineage was then subset and further annotated into subtypes by integrating canonical markers and cluster-specific marker genes. Gene ontology (GO) enrichment analysis was performed using clusterProfiler[33], in which differentially expressed genes between two conditions of interest were identified with sc.tl.rank_genes_groups with the “wilcoxon” method, filtered by FDR<0.05 and |logFC|>0.5 or 1, and subsequently subjected to enrichment analysis.

### Construction of the immune microenvironment and cell type enrichment analysis

The correlation matrix (Fig. 1D and 2A) describing the immune microenvironment was constructed by first calculating the proportion of each cell type in each patient, followed by computing Spearman correlation coefficients between cell types across all patients (Fig. 1D) or CSF patients only (Fig. 2A) using these proportions as features.

For modeling the relationship between cell proportions and disease status, age and sex were included as covariates. Generalized linear model (GLM) regression was performed using the GLM function from statsmodels.formula.api[34] with the formula: cell proportion ∼ disease status + age + sex, with disease status using “HC” as the reference, age using “<30 years” as the reference, and sex using “female” as the reference. Ro/e [35] was defined as the observed proportion of a given cell type in a specific group divided by its overall proportion across all patients. To account for the influence of outliers in the cell composition analysis, we applied an interquartile range (IQR)-based filtering procedure to remove upper-tail outliers. For each cell type in a given patient group, we first computed its (Q1) and third (Q3) quartiles of all non-zero proportions for that cell type across all patients in that patient group. Then the IQR was defined as Q3 - Q1, and the upper threshold was set to Q3 + 1.5 × IQR. Patients with values exceeding this threshold were considered outliers and excluded. Outlier detection was performed separately for each cell type in each patient group defined by different combinations of disease status, age group, and sex. Groups with fewer than three patients were exempted from outlier filtering.

### Interaction analysis

Cell-cell interaction analysis was performed using CellChat[36]. Based on the ligand-receptor interaction database CellChatDB, the computeCommunProb function was used to calculate the probability of specific ligand-receptor interactions between pairs of cell populations. The computeCommunProbPathway function was used to infer interactions at the signaling pathway level and to analyze the functional roles of each cell population within the pathway. The netVisual_bubble function was used to visualize the interaction probabilities of specific ligand-receptor pairs between cell types. The rankNet function was used to compare the significance of pathway strength or number between two datasets, with p=0.05 as the threshold.

### VDJ analysis and trajectory inference

VDJ analysis was conducted using Scirpy 1.20[37], and STARTRAC[35] was employed to calculate the migration capacity (migr) of each cell type across tissues and the transition potential (tran) between cell types. To construct the cell trajectory network for a given patient set, STARTRAC was first used to estimate the transition potential of T cell types in that patient set. For each pair of cell types, if the tran value was nonzero in at least three patients, these cell types were considered to undergo transition, with the degree of transition represented by the mean tran value across all patients.

## Discussions

The pathogenesis of NPSLE remains incompletely understood but is believed to involve complex inflammatory and vascular mechanisms, with blood-brain barrier (BBB) disruption playing a central role. BBB dysfunction facilitates the entry of various pathological factors into the CNS[4,5], thereby exacerbating neuronal damage through the actions of autoantibodies, cytokines, and immune cells[6]. Accumulating evidence demonstrates elevated levels of inflammatory cytokines (e.g., IL-6, IFN-α, TNF-α)[7,8], and autoantibodies (e.g., anti-ribosomal P, anti-NR2, antiphospholipid antibodies)[9–11] in the CSF of NPSLE patients, underscoring the significant roles of inflammation and immune activation in disease progression. Over the past decades, CSF single-cell studies have been increasingly applied to identify disease-specific immune cell states, cytokine profiles, and potential biomarkers in various CNS diseases, such as Lewy body dementia[12], multiple sclerosis[13] and Alzheimer’ s disease[14], revealing novel and highly reliable pathological mechanisms and potential clinical applications.

In this study, we performed, for the first time, single-cell transcriptomic profiling of CSF and paired blood samples from patients with NPSLE, providing an unprecedented view of the immune landscape in this condition. By integrating single-cell RNA sequencing with single-cell VDJ sequencing, we constructed a comprehensive CSF–blood single-cell atlas that delineates the immune microenvironmental differences across tissues and among categories of CNS diseases. Our results also indicate that both innate and adaptive immune responses jointly contribute to the pathogenesis of NPSLE. We identified BAM-CCL3 as a population exhibiting strong recruitment capacity and found that, compared to HCs, BAM-CCL3 was enriched in the CSF of NPSLE patients, which may be closely associated with the inflammatory response. We also observed a strong activation of Bm cells and CD4^+^ Tregs in the CSF of NPSLE patients. Moreover, CD8^+^ Tem cells in NPSLE exhibited migration from the blood into the CSF, accompanied by notable clonal expansion. These results indicate the presence of persistent CNS-localized adaptive immune dysregulation. Conceivably, some, if not all, NPSLE patients may exhibit a strong immune-inflammatory phenotype, suggesting the potential for precise immunotherapeutic interventions targeting B cell and T cell pathways. In addition, we established the scCDCB database (https://sccdcb.gao-lab.org) as a reference framework for future studies on CNS diseases. These findings offer novel insights into the immunopathogenesis of NPSLE and highlight potential cellular and molecular targets for future therapeutic development.

In our study, we reported for the first time a marked enrichment of BAM-CCL3, a chemokine-expressing subset of BAMs, in the CSF of patients with NPSLE. Residing at the boundaries of the CNS, BAMs can regulate immune cell entry into the meningeal spaces and neuroinflammatory responses [54]. Accordingly, the even stronger expression of chemokines by BAM-CCL3 in NPSLE is likely to recruit pathogenic T cells into the brain in CNS diseases [55], inducing immune imbalance and thus NPSLE pathogenesis. In particular, our data showed that one of the most active chemokine axes here is CXCL16-CXCR6, which is associated with the pathogenesis and disease activity of not only multiple autoimmune diseases including multiple sclerosis, autoimmune hepatitis, rheumatoid arthritis, Crohn’s disease, and psoriasis [56], but also between microglia and CD8^+^ T cells in Alzheimer’s disease [57]. Conceivably, CXCL16-CXCR6 has been considered as a potent drug target in CNS diseases[58], with effective blockade CXCL16 [59] or CXCR6 [60] demonstrated in animal models of experimental autoimmune encephalomyelitis. We also note that BAM-CCL3 might also mediate NPSLE endothelial activation and subsequent barrier injury by upregulating VEGF family genes; in line with this, Pedragosa et al. also demonstrated BAMs’proinflammatory role in neuroinflammation after ischemic stroke by partly through increasing cortically secreted VEGF protein expression[61]. Taken together, BAM-CCL3 may be a prospective target for further investigation.

Our findings on elevated BAFF and APRIL signaling with memory B cells in NPSLE CSF align with previous human studies [62]. Because BAFF and APRIL belong to the TNF family of cytokines and promote B cell activation and differentiation [22], it is expected that NPSLE patients have more immunoglobulins, possibly locally produced, in CSF than HCs. Indeed, in NPSLE patients, oligoclonal bands of immunoglobulins were detectable in 26.5% of CSF samples, and elevated intrathecal IgG indices were also observed [63], together with reports of anti-LGI1 antibodies-associated NPSLE[64], supporting the involvement of adaptive immune mechanisms driven by B cells within the CNS. Notably, this mechanism were also observed in other CNS diseases, such as neuromyelitis optica spectrum disorder (NMOSD) [65], anti-N-methyl-D-aspartate receptor antibody (anti-NMDAR) encephalitis [66], and MS [67], suggesting a CNS-related disease pathology that exacerbates CNS inflammation. Although B-cell-depleting agents are increasingly included in the management of NPSLE in the most recent ACR SLE guideline [68], evidence from randomized trials remains limited. Observational data suggest that refractory cases might benefit from rituximab[69,70,71]. However, systemic anti-CD20 monoclonal antibodies, while revolutionizing B-cell – targeted therapy, appear to have limited ability to penetrate the BBB when administered intravenously, meaning that CNS B-cell reservoirs may be spared or cleared only slowly [72,73]. Consequently, newer therapeutic approaches that are confirmed or hypothesized to better access the CNS — for example, CD19-CAR T-cell therapy — might hold greater promise for depleting intrathecal or meningeal B-cell populations and achieving more effective and durable immunomodulation in neuro-immunological disorders [74–77].

Brain-resident Tregs expand in multiple neuroinflammatory pathologies, including MS and traumatic brain injury [78]. In this study, Treg cells in NPSLE CSF exhibited enrichment and activation despite prominent inflammation, possibly favoring a long-term persistence within the CSF and suppression of activated immune cells. Of note, similar results were also observed in the CSF of MS patients, where Tregs accumulate but only exert limited control over glial cell-driven chronic injury and thus cannot fully prevent neurodegeneration [79], and other autoimmune diseases, where Tregs are enriched in inflamed tissues, but do not effectively suppress autoimmune responses[80,81], suggesting a pan-disease role of Treg-based limited immunosuppression in these diseases. Mechanistically, Korn et al. suggested that Treg immunosuppressive function is context dependent, influenced by the inflammatory microenvironment [82], providing a possible future direction of studying whether and how Treg enrichment in NPSLE CSF fails to compensate for CNS inflammation and neural injury.

We observed that clonal expansion in NPSLE CSF was mostly observed in CD8^+^ Tem, with a potential preference for migration from blood into CSF, suggesting its functional role in NPSLE pathology. Similar results of CD8^+^ T cells in other CNS diseases were discovered by previous studies, where these cells were shown to potentially contribute to tissue damage in inflammatory demyelination animal models[83,84], to CNS tissue damage in MS lesions[84], and to the pathogenesis of EAE[85]. We note, however, that in healthy (old in particular) individuals, activated T cells and especially activated CD8^+^ T subsets, also frequently exhibit high level of clonal expansions[86,87], possibly reflecting ongoing physiological surveillance by the CNS[88,89]. Therefore, relevant factors such as age must be considered when further examining the role of clonally expanded CD8+ T cells with an effector/memory phenotype in NPSLE pathology.

We also preliminarily investigated whether there is any potential dysregulation of CD8^+^ T differentiation in NPSLE. At baseline analysis in which NPSLE and HC were mixed together, we found that CD8^+^ T cells in PBMCs preferentially differentiate along the trajectories between CD8-Trm-KLRG1 and CD8-Trm-ZNF683, and from CD8-Tem to CD8-Temra-GNLY and CD8-Temra-CMC1 (Fig. S5A), while in CSF, CD8^+^ T cells preferentially differentiated from many CD8^+^ T subsets toward CD8-Tex, as well as between CD8-Temra-GNLY and CD8-Temra-CMC1 but not from CD8Tem to CD8Temra, indicating a stronger tendency toward exhaustion in the CSF (Fig. S5B). When NPSLE and HC blood or CSF samples were analyzed separately by constructing differentiation trajectories for each group, we identified a subtle CSF difference in exhaustion progression (Fig. S5C, D) but no major differences in the CD8Tem and CD8Temra transition in peripheral blood (Fig. S5E, F). Specifically, in NPSLE CSF, both CD8-Temra-GNLY and CD8-Trm-ZNF683 exhibit trajectories toward exhaustion (Fig. S5F), whereas in HC, exhaustion is mainly driven by CD8-Tex_pre and CD8-Teff (Fig. S5D). Since CD8Tex cells are predominantly enriched in the CSF and largely absent from PBMCs, the differences above are more likely derived from local differentiation rather than from peripheral infiltration across the BBB, suggesting that in NPSLE, within-CSF microenvironment factors such as persistent antigenic stimulation might hamper the proper development -- and drive the exhaustion -- of specific CD8⁺ T-cell subsets that exert their functions normally in HC.

Of interest, we noticed pronounced different immune microenvironments between CSF and blood (Fig. S6A). In addition to statistically significant differences in proportions (Fig. S6B), we also found that multiple cells including border-associated macrophages(BAMs), central memory T(Tcm) cells, effector memory T(Tem) cells, effector T(Teff) cells, tissue-resident memory T(Trm) cells, exhausted T(Tex) cells, and regulatory T(Treg) cells, are predominantly enriched in CSF, whereas other cell types like memory B(Bm) cells, naïve B(Bn) cells, naïve T(Tn) cells, and terminally differentiated effector memory T(Temra) cells are enriched in PBMCs. Specifically, while blood T cells are primarily naïve and terminally differentiated phenotypes, their CSF counterparts predominantly exhibit memory or effector phenotypes, consistent with previous studies[90] and indicating that T cells in CSF largely represent antigen-experienced populations that have undergone differentiation, rather than naïve thymic emigrants directly entering the CSF. In particular, the CSF enrichment in Tcm cells and Tem cells, two cell types characterized by long time immunological memory and rapid recall responses, reflects a CSF immune landscape that is more oriented toward immune surveillance and regulation.

Admittedly, this study has several limitations. Firstly, the number of NPSLE patients with paired blood-CSF remains limited, underscoring the need to expand sample size in future studies to not only derive cellular, pathway, and TCR-based conclusions that are more robust against confounding factors such as sex and age, but also reveal key cellular and molecular differences among NPSLE subtypes, including seizures and cerebrovascular disease. In addition, the present study is based solely on transcriptomics and TCR repertoire, and the corresponding results would benefit from additional omics for validation and interpretation, including, but not limited to, epigenetics (e.g. single-cell chromatin accessibility), BCR repertoire, and proteomics, as well as functional experimental assays. Last but not least, although CSF is widely used for diagnosing CNS diseases, it does not capture most neuronal cells and their microenvironments at the lesion that reflects the root disease mechanisms; therefore, brain parenchyma samples from (e.g. postmortem) NPSLE patients, provided that they are appropriate ethical and clinical criteria, would greatly advance the understanding of NPSLE pathology and its development of therapeutics.

## Data and code availability

The raw sequence data reported in this paper are available in the Genome Sequence Archive[91] in National Genomics Data Center[92], China National Center for Bioinformation / Beijing Institute of Genomics, Chinese Academy of Sciences (GSA-Human: HRA010161) that are publicly accessible at https://ngdc.cncb.ac.cn/gsa-human. Requests for additional materials can be made via email to the corresponding authors.

All previously published single-cell datasets used for building the scCDCB are available from their publications (Table 1). All processed data can be accessed at https://sccdcb.gao-lab.org.

All scripts used for data analysis, including the processing of the atlas, as well as the generation of all results in figures, are available from GitHub at https://github.com/gao-lab/scCDCB.

## Competing interests

The authors declare that they have no competing interests.

## Acknowledgements

This study was supported by the Chinese National Key Technology R&D Program, Ministry of Science and Technology (2021YFC2501300), National Natural Science Foundation of China (32441090, 92574108), CAMS Innovation Fund for Medical Sciences (CIFMS) (2021-I2M-1-005, 2022-I2M-1-004, 2023-I2M-2-005, 2025-I2M-XHZY-001, XHJC-005), National High Level Hospital Clinical Research Funding (2025-PUMCH-D-001, C-010, C-026, C-054, 2022-PUMCH-D-009), Beijing High-Level Innovation and Entrepreneurship Talent Support Program (G202511002, G202521084), Peking Union Medical College Hospital Talent Cultivation Program (B-UGG10647), and the National Science and Technology Major Project (2022ZD0115004). Part of the analysis was carried out on the Computing Platform of the Center for Life Sciences of Peking University and supported by the High-performance Computing Platform of Peking University and Changping Laboratory (2021D-01-01-04).

**Sup Fig. 1 Quality metric distribution of single-cell RNA and VDJ data from blood and CSF.**

**A, B**. Quality metric distribution of CSF (A) and blood (B) single-cell RNA data for each study. **C, D.** Overall quality metric distribution of CSF (C) and blood (D) single-cell RNA data, including gene counts, total counts, and the proportion of mitochondrial gene expression. **E, F.** Distribution of gene counts per cell in CSF (E) and blood (F) single-cell transcriptome data. Red lines indicate the upper and lower thresholds of gene count used for filtering low-quality cells, with the lower threshold set at 700 and the upper threshold at 6000. **G.** Distribution of chain pairing results for single-cell VDJ data. H. UMAP plot showing the studies.

**Sup Fig. 2 Expression of subtype markers.**

**A-E.** The expression of markers for CD8^+^ T cells, CD4^+^ T cells, NK cells, B cells, and myeloid cells, respectively.

**Sup Fig. 3 Signaling Differences Between NPSLE and HCs**

**A.** Differences in information flow of each signaling pathway between NPSLE and HC CSF, with BAM-CCL3 acting as the signal sender. **B,C.** Differences in information flow of each signaling pathway between NPSLE and HC, with Bm-IGLC2 and Bm-IGKC combined as the signal receiver (B) or signal sender (C). **D. D**istribution of communication probabilities of the CD70-CD27 axis sent out by Bm-IGLC2 and Bm-IGKC, filtered by p<0.05. E. Differences in information flow of each signaling pathway between NPSLE and HC, with CD4Treg-CCR7 and CD4Treg-HLA combined as the signal receiver.

**Sup Fig. 4 Clonal Expansion of T Cells.**

**A.** The top 30 expanded clone IDs in the CSF cells and PBMCs of NPSLE and HC, standardized by the number of cells in each patient. **B,C.** The distribution of different tissues (B) and diseases (or HC; C) in each expanded clone. Each point represents a clone ID, and the size indicates the extent of clonal expansion. **D-G**, The distribution of cell types in each expanded clone, in HC CSF (D), NPSLE CSF (E), HC blood (F), or NPSLE blood (G).

**Sup Fig. 5 CD8 T cell differentiation network.**

**A,B.** CD8^+^ T differentiation trajectories computed from STARTRAC-based transition values within CSF cells (A) or PBMCs (B). Cell-type pairs with non-zero transition values in at least three patients were retained, and the mean transition value was computed across these patients. For both the heatmap and the network visualization, the displayed values represent the normalized mean transition values across all patients. **C-F.** CD8^+^ T cell transition networks and the normalized mean transition value heatmaps within HC PBMCs (C), HC CSF cells (D), NPSLE PBMCs (E), and NPSLE CSF cells (F), respectively.

**Sup Fig. 6 Distinct Immune Microenvironmental Composition Between Blood and CSF.**

**A.** The heatmap of the Spearman’s correlation coefficients (ρ) on cell type proportions between different patients, with the top two color bars showing the distribution of the ratio of enrichment (Ro/e) for each cell type across different tissues. Cell type colors follow those in panel C. B. Paired Wilcoxon rank-sum tests using the cell type proportions from paired blood and CSF samples. Orange x-axis labels represent an enrichment in CSF, and blue represents an enrichment in blood.

**Sup Table S1. CSF and blood samples infomation of published data.**

**Sup Table S2. Clinical, immunological, and treatment characteristics of NPSLE patients at the time of sample collection.**

**Sup Table S3. DEGs in NPSLE CSF cells relative to HC CSF cells.**

## Notes

### Competing Interest Statement

The authors have declared no competing interest.

