## Supplementary figures and images for "CNS diseases cerebrospinal fluid single-cell atlas reveals immune characteristics of neuropsychiatric systemic lupus erythematosus"

### Supplementary Figures

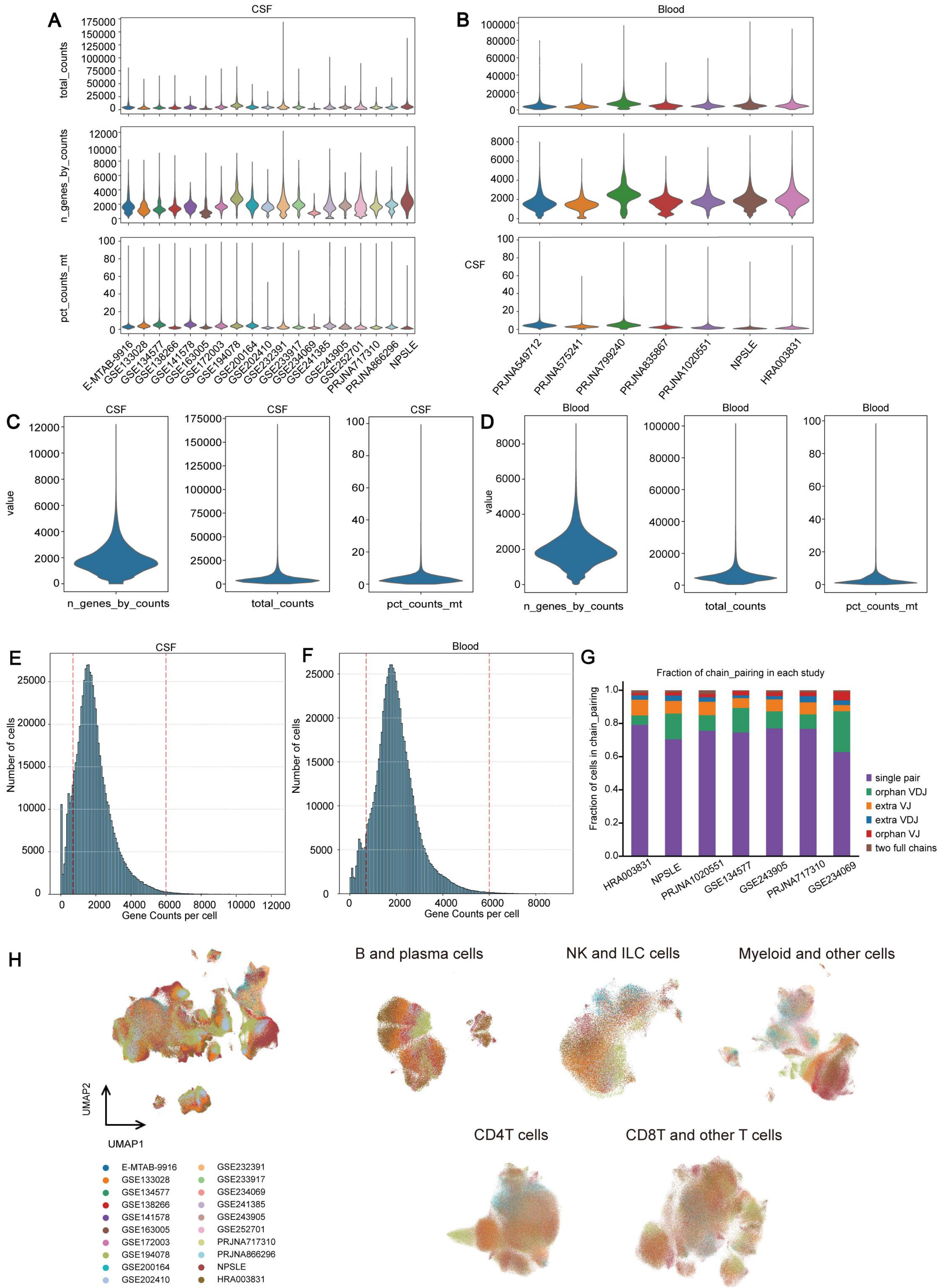

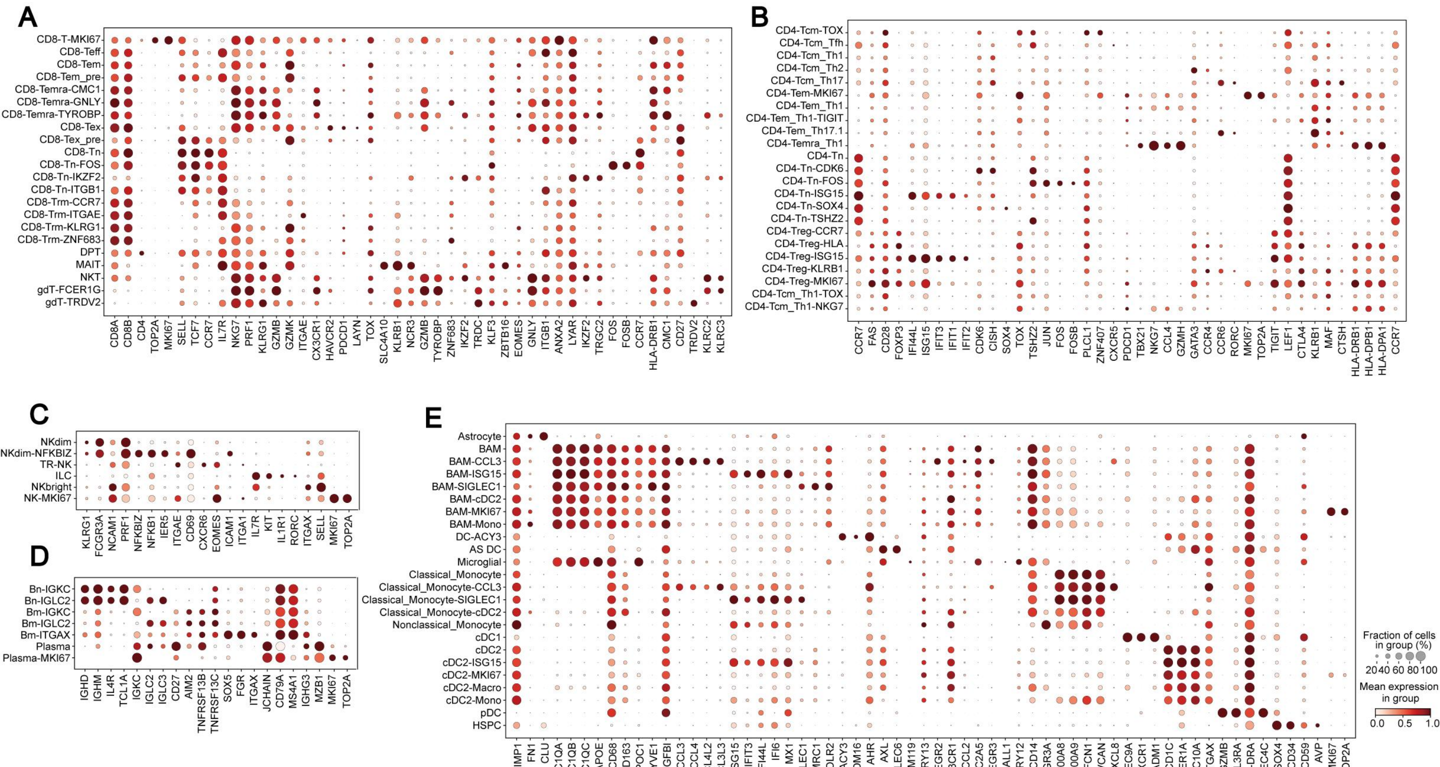

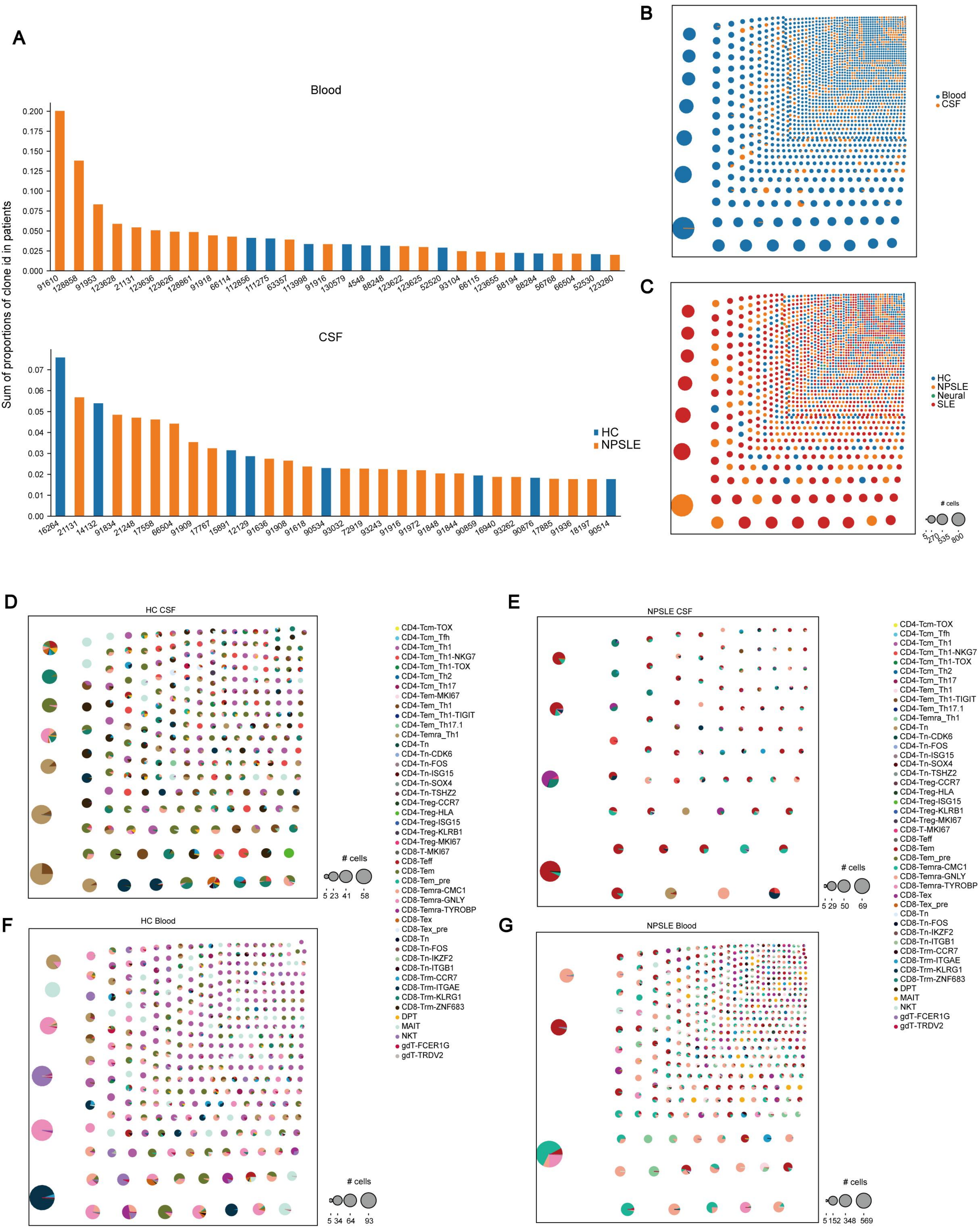

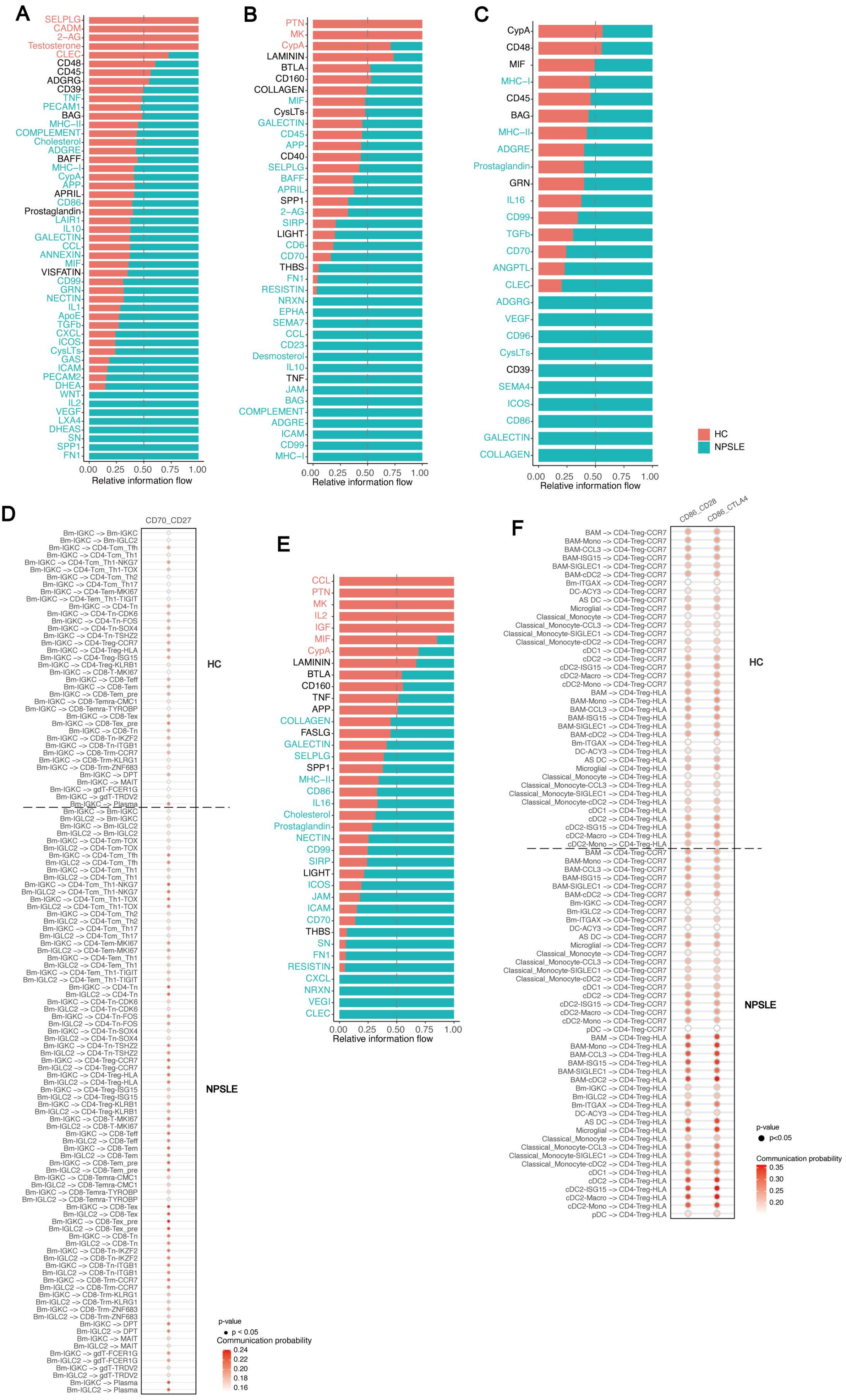

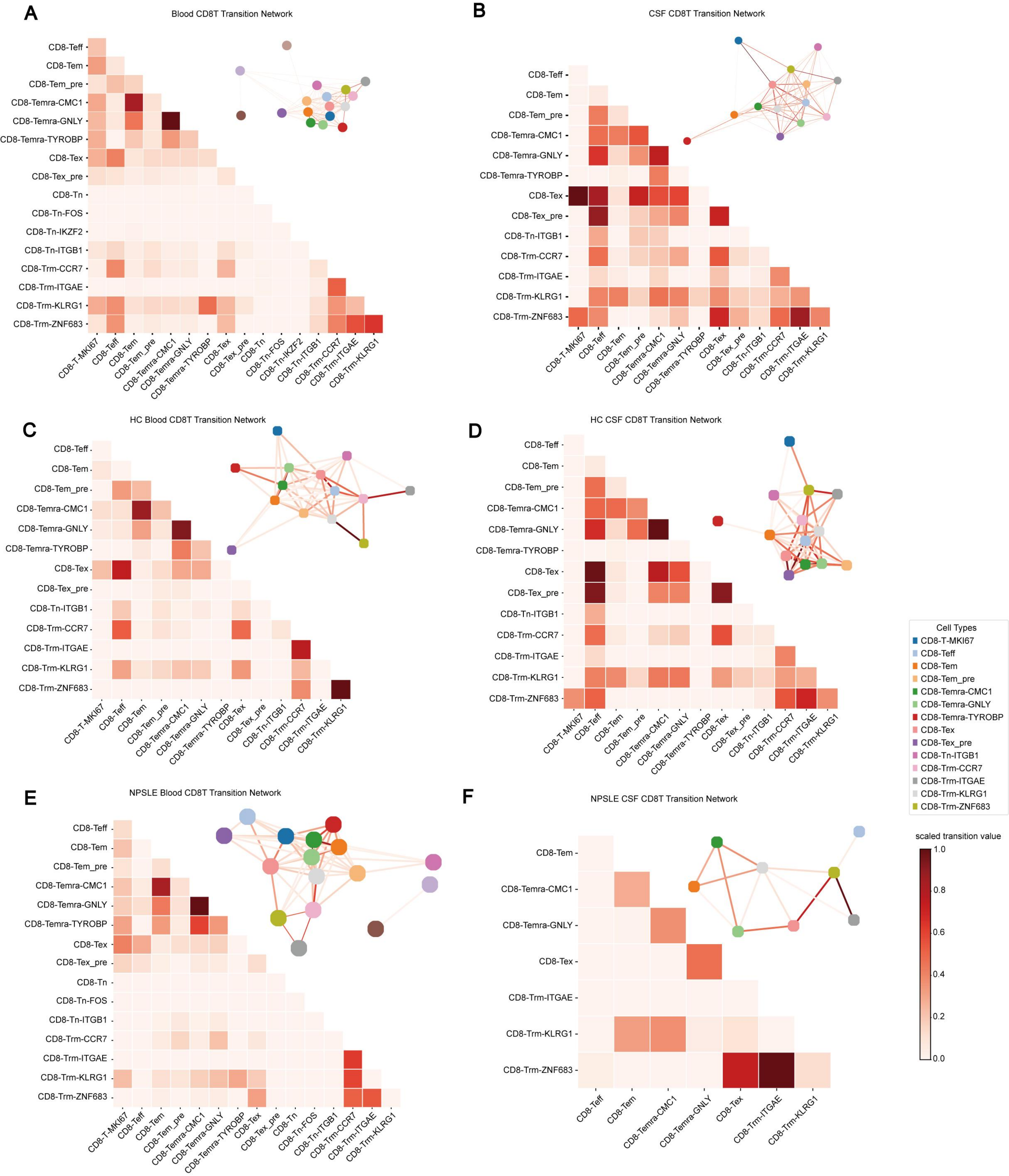

Sup Fig. 6

A

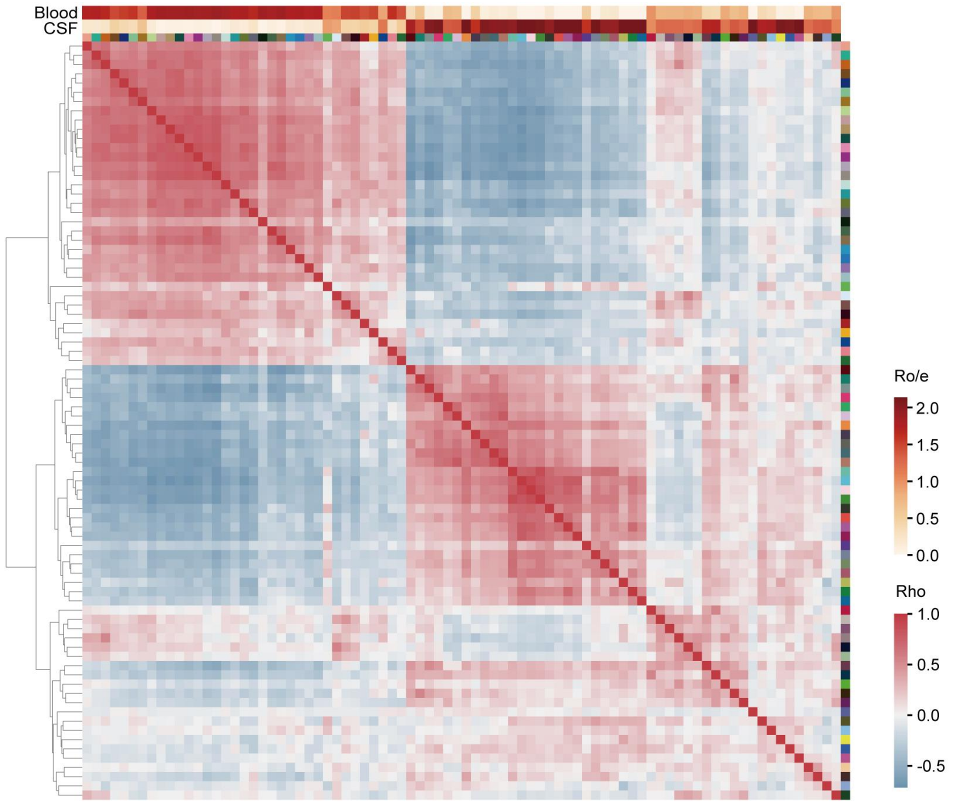


B

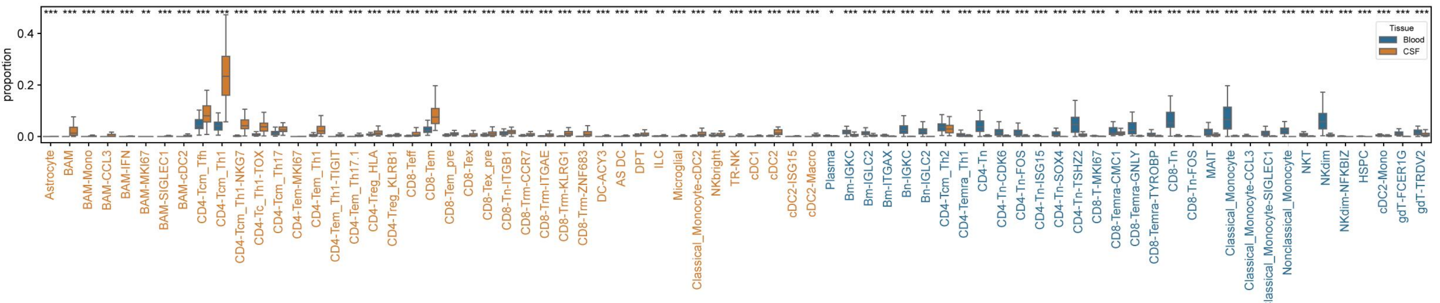


* 0.01<FDR<0.05
** 0.001<FDR<0.01
*** FDR<0.001
