## Supplementary Table 2 for "CNS diseases cerebrospinal fluid single-cell atlas reveals immune characteristics of neuropsychiatric systemic lupus erythematosus"

**Supplementary Table S2. Clinical, immunological, and treatment characteristics of NPSLE patients at the time of sample collection**

|  | Sex | Age |  | SLE disease duration | NPSLE duration | Extra-NP organ involvement | C3 level | C4  Level | Anti-dsDNA | aPL | Treatment |
| --- | --- | --- | --- | --- | --- | --- | --- | --- | --- | --- | --- |
| Case1 | F | 22y | Seizure | 3 weeks | 1 week | Renal, hematological, mucocutaneous, and pulmonary involvement | 0.245 | 0.015 | Positive | Positive | Steroid pulse therapy, HCQ |
| Case2 | F | 17y | Myelopathy | 3 weeks | 1 week | Hematological involvement; myositis | 0.337 | 0.015 | Positive | Negative | Steroid pulse therapy, HCQ |
| Case3 | F | 36y | Myelopathy | 11 years | 5 years | Arthritis | 0.327 | 0.033 | Positive | Positive | Steroids, MMF, HCQ |
| Case4 | F | 43y | CVD | 2 years | 2 years | Renal, hematological, and pulmonary involvement | 1.034 | 0.220 | Positive | Positive | Steroids, AZA，HCQ |
| Case5 | F | 54y | CVD | 7 years | 1 month | Hematological involvement | 0.844 | 0.129 | Negative | Negative | Steroids, MMF, HCQ, |
| Case6 | M | 38y | Seizure | 4 months | 4 months | Renal involvement | 0.363 | 0.005 | Positive | Positive | - |
| Case7 | F | 44y | ACS | 18 years | 2 weeks | Renal, hematological， and gastrointestinal involvement | 0.509 | 0.33 | Positive | Negative | Steroids |
| Case8 | F | 48y | Seizure | 21 years | 20 years | Renal and mucocutaneous involvement | 0.501 | 0.030 | Positive | Negative | Steroids, MMF, HCQ |
| Case9 | F | 30y | Seizure | 2 years | 2 years | Renal involvement | 0.616 | 0.125 | Positive | Negative | Steroids, HCQ |

ACS: acute confusional state; aPL: antiphospholipid antibody; anti-dsDNA: anti-double-stranded deoxyribonucleic acid antibody; AZA: azathioprine; C3: complement component 3; C4: complement component 4; CVD: cerebrovascular disease; F: female; HCQ: hydroxychloroquine; M; male; MMF: mycophenolate mofetil; NP: neuropsychiatric; NPSLE: neuropsychiatric systemic lupus erythematosus; SLE: systemic lupus erythematosus.
